# Cortical vestibular-evoked potentials depend on body orientation

**DOI:** 10.64898/2026.03.10.705553

**Authors:** Paul Kobliska, Lou Seropian, Yannick Becker, Yves Cazals, Christophe Lopez

## Abstract

Otolithic vestibular receptors encode linear acceleration and head orientation relative to gravity, providing a fundamental reference signal for perception, action, and higher-order cognitive functions. However, the cerebral dynamics of otolithic information processing and their sensitivity to body orientation remain poorly characterized. Here, we used sound-induced vestibular stimulation combined with electroencephalography (EEG) in human participants to characterize vestibular-evoked potentials (vEPs) and to examine how body orientation relative to gravity modulates these responses. Otolithic activation was validated using cervical vestibular-evoked myogenic potentials, confirming activation of otolithic pathways by 105 dB and 500 Hz tone pips. Acoustic stimuli included a novel masked stimulus that reduced auditory perception while preserving vestibular activation. Sound-induced vestibular stimulation elicited reliable short- and middle-latency vEP components. Importantly, middle-latency components Na/Pa (peak latency: 20–30 ms) and N*/P* (41–54 ms) were modulated by body orientation, showing reduced amplitudes in the supine compared with the upright position, but only for otolithic-activating sounds and not for matched auditory control stimuli. This orientation-dependent modulation was stronger for Na/Pa and was specific to otolithic-activating sounds, supporting a vestibular rather than auditory origin and highlighting early integration processes. This pattern suggests that modulation of vEPs primarily reflects context-dependent cerebral processing of otolithic signals rather than peripheral sensory mechanisms. Together, our results establish Na/Pa and N*/P* as reliable cerebral markers of otolithic processing and demonstrate that early cerebral vestibular responses are dynamically shaped by body orientation and postural context.

**NEW & NOTEWORTHY:** This study identifies middle-latency vestibular-evoked potentials as reliable cerebral markers of otolithic information processing. We show that both Na/Pa and N*/P* are selectively modulated by body orientation for otolithic-activating sounds, but not for auditory controls. These findings reveal that early otolithic vestibular processing dynamically depends on bodily context, extending current models of multisensory vestibular integration.

## INTRODUCTION

Vestibular otolithic receptors encode linear translations (including gravitational acceleration) and head tilts relative to gravity. These signals are essential for reflexive control of posture, gait, and gaze stabilization (1). Beyond these reflex functions, otolithic information, which is closely tied to gravity, is crucial for perceiving verticality, distinguishing the direction of “up” and “down” (2), and constructing internal models of visual object motion in gravitational fields (3). Otolithic signals also contribute to a range of higher-level cognitive processes, including visual perception of bodies (4), spatial navigation and memory (5), own-body perception (6), and self-consciousness (7).

The role of otolithic information in these diverse sensorimotor and cognitive functions is supported by widespread vestibular projections to the cerebellum, thalamus, and a large network of cortical areas (8–12). However, understanding the neural bases of otolithic information processing remains limited, mainly due to methodological constraints in neuroimaging, which prevent performing naturalistic body translations in MRI or PET scanners. Interestingly, mammalian otolithic receptors are also sensitive to high-intensity acoustic stimulation (13–16), which is thought to be an evolutionary remnant. Electrophysiological studies in rodents showed that otolithic receptors respond to specific intensity (around 100 dB SPL) and frequency (500 Hz) sounds, such as short tone pips (16, 17). Such stimuli can generate fluid vibrations within the inner ear, directly activating hair bundles from otolithic mechanoreceptors (17, 18). This stimulation method, here referred to as sound-induced vestibular stimulation (SVS), offers a well-controlled, non-invasive approach to studying otolithic vestibular pathways and is compatible with functional MRI (19), non-invasive electroencephalography (EEG) (20), and invasive stereo-EEG (21, 22).

SVS is widely used in clinical settings to assess vestibular-evoked myogenic potentials (VEMPs) via electromyography (EMG) (23). Cervical VEMPs (cVEMPs), recorded from the sternocleidomastoid muscles, reflect vestibulo-collic reflex pathways and primarily indicate saccular activation, characterized by a biphasic P13/N23 response (16, 24–26). Ocular VEMPs (oVEMPs), recorded from inferior oblique eye muscle, reflect vestibulo-ocular reflex pathways, primarily indicating utricular activation, and are characterized by a biphasic N10/P22 response (23, 27, 28).

Data on brain responses to SVS are limited. Functional MRI studies using SVS have shown activation in the parieto-temporal cortex, superior temporal gyrus, posterior insula, parietal operculum (29–32), as well as in the prefrontal cortex, premotor cortex, frontal eye fields, and cingulate cortex (19). A meta-analysis of functional neuroimaging data localized otolithic responses primarily in the parietal operculum/retroinsular cortex and superior temporal gyrus (33). However, as the low temporal resolution of functional MRI limits our understanding of the spatiotemporal dynamics of otolithic information processing, developing electroencephalographic descriptions of vestibular-evoked potentials (vEPs) seems necessary (reviewed in (34)).

Cerebral vEPs using SVS were first observed in 1980 in guinea pigs (14) and adapted for human studies in the early 2000s to 2010s (35, 36, 20). Human EEG studies employing SVS have linked vestibular processing to short-latency biphasic P10/N17 components over parietal (20) and parieto-occipital electrodes (35, 37), and to a frontal N15/P21 component. Several studies proposed that middle-latency components, notably the biphasic N42/P52 (also noted N*/P*) recorded at the FCz electrode, are also key markers of otolithic information processing (20). Late-latency parieto-temporal components, such as P60, N70, and P110, may also indicate vestibular activity (22). While cochlear activation by SVS could suggest a mixed vestibular-cochlear origin for vEPs, the presence of short-latency components in individuals with bilateral hearing loss (35, 37), and absence of middle-latency components in patients with unilateral vestibular loss (20), indicate that middle-latency components are relevant markers of vestibular processing.

This study aimed to refine otolithic responses characterization in the human brain using EEG and to examine the influence of body orientation on these responses. Previous vEPs studies have primarily investigated the effects of acoustic stimuli varying in intensity and frequency, but otolithic receptors also encode gravitational acceleration, generating distinct signals based on head orientation. There is robust experimental evidence that otolithic afferent discharge frequency is modulated by static head orientation with respect to gravity (38–40) and orientation-dependent vestibular responses have been recorded in multiple brain regions, including the cerebellum, thalamus and parieto-insular vestibular cortex (41–45). To explore these effects, we developed new acoustic stimuli targeting otolithic receptors and contrasted vEPs induced by these sounds with control sounds, systematically varying stimulus intensity, frequency, and morphology. Additionally, we compared vEPs recorded in upright sitting and supine positions, hypothesizing that head orientation modulates vEPs amplitude, particularly in middle-latency components, thereby influencing vestibulo-thalamo-cortical processing.

## MATERIALS AND METHODS

### 1. Participants

Twenty participants with no history of otoneurological disorder and hearing impairment were initially recruited in this study. Two participants were excluded from the EEG analyses due to the absence of cVEMPs, resulting in a final sample of 18 participants (10 women; mean age ± SD: 24.5 ± 4.3 years). All participants were right-handed according to the Edinburgh Handedness Inventory (all scores ≧ 40%) (46). The study protocol was approved by the Research Ethics Committee and participants gave their written informed consent to participate in the study.

### 2. Acoustic stimuli

Four acoustic stimuli were designed to activate the otolithic system or to control for sound intensity and frequency (**Figure 1a**):

- *Activating sound*. The first acoustic stimulus designed to evoke vestibular responses consisted of a 500 Hz air-conducted tone pip of 5 ms (rise and fall time: 2 ms, plateau: 1 ms) presented at 105 dB SPL known to activate otolithic receptors (16, 17, 20, 24, 47).
- *Intensity control*. To control for sound intensity, a tone pip with an identical structure and frequency (500 Hz) as the *Activating sound* was presented at 60 dB SPL. This stimulus at intensity below the vestibular threshold should not evoke any otolithic response (20, 22).
- *Frequency control*. To control for sound frequency, a tone pip with an identical structure and same intensity (105 dB SPL) as the *Activating sound* was presented at a frequency of 2500 Hz. This stimulus should not evoke any otolithic response (22, 48).
- *Masked activating sound*. A new stimulus was designed to acoustically mask the activating tone pip. The *Activating sound* was covered by a 90 dB SPL envelope of noise with a frequency ranging from 125 to 2000 Hz. This envelope of noise was added from 50 ms before to 50 ms after the tone pip onset. This stimulus should evoke a vestibular response, while masking the auditory component of the activating tone pip perception (49, 50).

**Figure 1.**
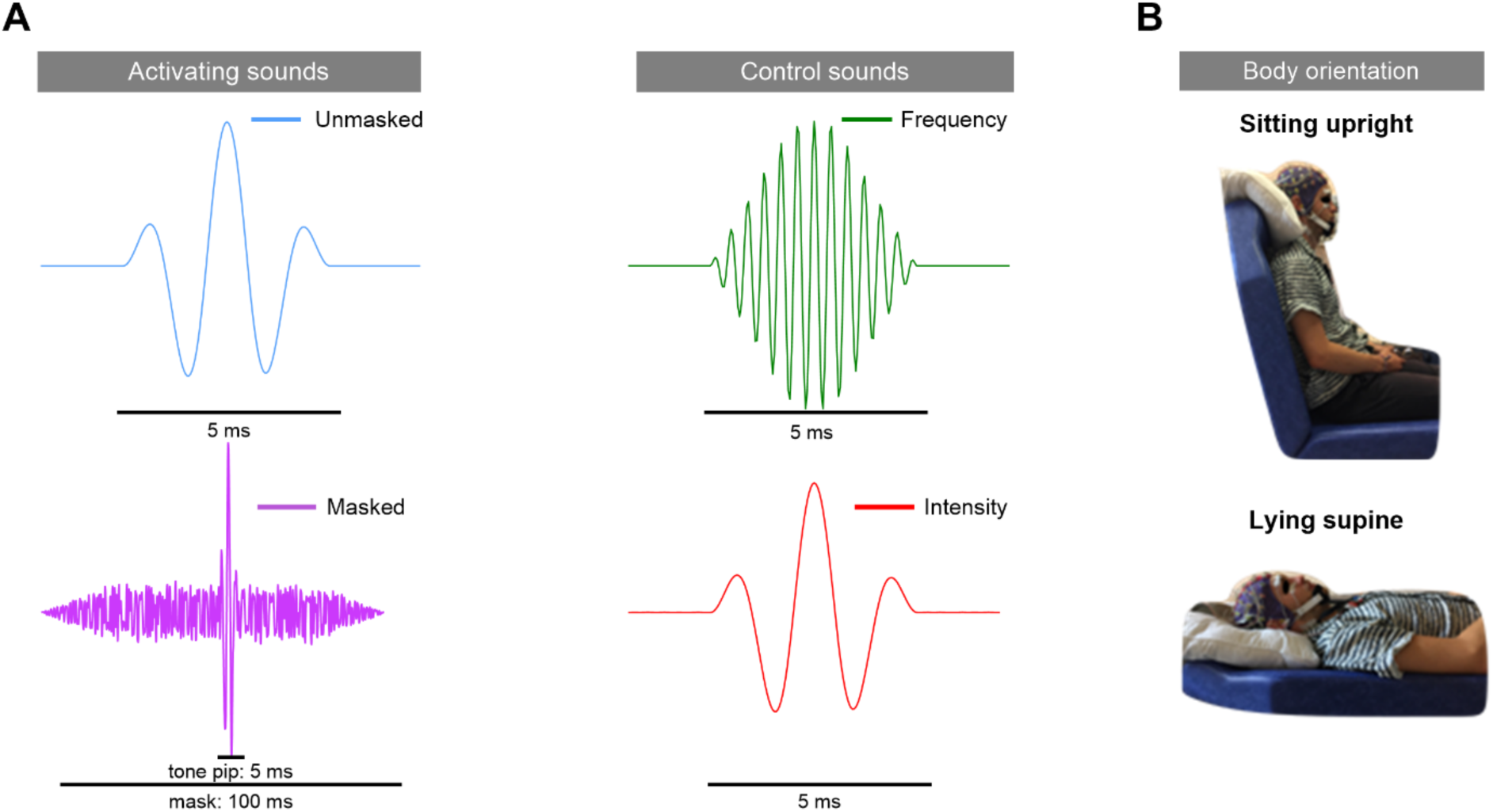
**(A)** Acoustic stimuli. The *Unmasked activating sound* was specifically designed to activate vestibular pathways and consists of a 500 Hz tone pip presented at 105 dB SPL. The *Frequency control* and *Intensity control* stimuli consist of a 2500 Hz tone pip at 105 dB SPL and a 500 Hz tone pip at 60 dB SPL, respectively. The *Masked activating sound* is a 500 Hz tone pip at 105 dB SPL, acoustically masked by a noise envelope at 90 dB SPL, with a frequency range of 125–2000 Hz, applied from 50 ms before to 50 ms after the tone pip onset. **(B)** Acoustic stimuli were presented to participants in two body orientations: sitting upright and lying supine

Acoustic stimuli were computed with Matlab R2015b (MathWorks Inc.) and delivered via a FIREFACE UCX (RME audio Inc.) audio interface. The set-up was calibrated prior to experimentation using an artificial ear (2CC coupler, Pistonphone 64 dB and 4131 microphone, Brüel and Kjær, Denmark). Stimuli were presented with intra-auricular earphones with disposable earplugs adapted to the diameter of the external acoustic meatus (ER3C, Etymotic Research Inc.). The right ear was stimulated in all participants to reduce hemispheric difference in audio-vestibular processing (20, 29).

### 3. Electrophysiological recordings

All electrophysiological signals were recorded with a Biosemi ActiveTwo system (Biosemi Inc., Amsterdam, the Netherlands) at 1024 Hz sample rate and were bandpass-filtered between 0.16 and 512 Hz.

#### 3.1 Cervical vestibular-evoked myogenic potentials

In the first part of the experiment, cVEMPs were recorded to assess otolithic receptors activation for each acoustic stimulus. Surface EMG was recorded from the sternocleidomastoid (SCM) muscles using Ag/AgCl electrodes. Active surface electrodes were placed over the middle of the SCM muscle belly, where the largest cVEMPs can be recorded (26), and were referenced to an electrode placed on the sternum. Participants were tested while seating on a chair with their back tilted 45° backwards from the vertical position. They were required to contract the right SCM (ipsilateral to the acoustic stimulation) by lifting their head forward and turning it to their left. Acoustic stimuli were presented while maintaining muscle contraction and fixating on a visual target at 2 m. For each acoustic stimulus, 200 repetitions were presented. Stimuli were presented in a randomized order in two blocks of 400 stimuli, with an interstimulus interval randomly selected between 300 and 400 ms. Participants rested between blocks of stimulation.

#### 3.2 Cerebral vestibular-evoked potentials

In the second part of the experiment, vEPs were recorded in two body orientations – sitting upright and lying supine (**Figure 1b**) – presented in a quasi-balanced order between participants. Continuous EEG was recorded from 64 active Ag/AgCl scalp electrodes mounted on a head cap fitting the participant’s head circumference (international 10−20 system). The Common Mode sense–Driven Right Leg (CMS–DRL) reference electrodes (specific ground electrodes forming a feedback loop with the participant) were located next to the parietal electrodes. Electrode offset was maintained below 20 µV.

Eye movements were monitored using an electrode on the skin above the eye ipsilateral to the acoustic stimulation (right supra-ocular electrode), and an electrode below the right eye (right infra-ocular electrode, rIO) and the left eye (left infra-ocular electrode, lIO) fixed with medical tape. Reference electrodes were placed on the earlobes.

In the sitting upright position, participants were required to maintain their gaze on a target 2 m in front of them, to relax their neck muscles, and avoid blinking and swallowing. Their head was supported by a headrest and a pillow. In the lying supine position, the back of the chair was tilted so as to be horizontal and participants were required to maintain their gaze on a target located 3 m on the ceiling, to relax their neck muscles, and avoid blinking and swallowing. Their head was also supported by a pillow. For each body orientation, 600 repetitions of each acoustic stimulus were presented in a randomized order in 6 blocks of 400 stimuli, with an inter-stimulus interval randomly selected between 300 and 400 ms. Participants rested between blocks of stimulation.

Signals from both the left and right SCM muscles, and from the right supra-ocular, rIO and lIO electrodes were recorded simultaneously with EEG to assess potential electromyogenic and electrooculographic contributions to the vEPs.

### 4. Data processing

Electrophysiological signals were preprocessed and analyzed using MNE-Python (51) and custom-made scripts in Python.

#### 4.1 Cervical vestibular-evoked myogenic potentials

Data were referenced to the sternum electrode and band-pass filtered between 1 and 200 Hz. Epochs were then set to 30 ms before to 200 ms after onset of the tone pip, with a baseline defined as the 30 ms pre-stimulation. Each epoch was baseline-corrected by subtracting the mean of the baseline period to each data point of the trial. Trials were averaged for each acoustic stimulus (*Activating sound, Intensity control, Frequency control, Masked activating sound*).

#### 4.2 Vestibular-evoked potentials

First, raw time courses and power spectral density of all channels were visually inspected. Channels showing abnormal activity were subsequently interpolated using spherical splines (52). On average, 5 ± 2 electrodes (mostly frontal and temporal) were interpolated for both upright and supine orientations. Data were then re-referenced to an average reference and band-pass filtered between 1 and 200 Hz. After manually rejecting data spans contaminated with movement artefacts or muscular activity, ocular artefacts such as blinks and saccades were removed using an independent component analysis (53). Epochs were defined from 50 ms before tone pip onset to 300 ms post-stimulus, with a baseline defined as the 50 ms pre-stimulation. A baseline correction was then applied to each epoch following the procedure described in *Section 4.1*. Trials for which the dynamic range exceeded 100 µV were automatically rejected, and selected trials were visually inspected to reject any other epoch with remaining artefacts. On average, 8 ± 5% and 6 ± 4% of trials were rejected for the upright and supine orientation, respectively. Stimulus-locked EPs were computed by averaging selected trials for each acoustic stimulus (*Activating sound*, *Intensity control*, *Frequency control*, *Masked activating sound*) and each body orientation (*Upright sitting*, *Lying supine*) separately. Signals recorded at right SCM muscle, referenced to the sternum, and under electrode lIO, referenced to the linked earlobes (20), corresponding to selected EEG trials were processed following the same procedures.

### 5. Statistical analyses

#### 5.1 Cluster-based permutation test

A spatiotemporal cluster-based permutation test (54), implemented in MNE-Python (51), was used to evaluate differences in potentials evoked by acoustic stimuli across different body orientations, focusing specifically on the interaction between Sound and Body orientation factors. The statistic test for the permutation analysis was derived from a two-way repeated-measures ANOVA. This analysis identifies clusters of individual samples based on their temporal and spatial adjacency. A cluster is defined as a group of samples in time and space whose *F*-values exceed a predefined cluster-forming threshold corresponding to a *p-*value < 0.05. The cluster-level statistic was calculated as the sum of the *F*-values for all samples within a cluster. To control for Type I errors (false positives), the data was subjected to 1,000 random permutations, where sample labels were shuffled between conditions. This generated a null distribution of the maximum cluster-level statistics that could occur by chance. The Monte Carlo estimate of the permutation *p*-value was calculated as the proportion of null distribution values exceeding the observed maximum cluster-level statistic in the experimental data. Two conditions were deemed significantly different if this *p*-value was below the critical alpha level (*α* = 0.05). This procedure was applied at the sensor level using data from 64 scalp electrodes, covering the time window from 0 to 70 ms after the onset of the acoustic stimulus, a period that was associated with short- and middle-latency vEPs (22, 35).

#### 5.2 Peak-to-peak amplitude analysis

##### 5.2.1 Cervical vestibular-evoked myogenic potentials

The peak-to-peak amplitudes of the biphasic P1/N1 response were measured over the right SCM muscle (ipsilateral to the acoustic stimuli) for each acoustic stimulus. The peak-to-peak amplitude for biphasic components (e.g., P1/N1) was calculated as the difference between the amplitudes of the two opposite peaks. A Friedman’s ANOVA was performed to evaluate the effect of the within-subject factor *Sound* (*Activating sound*, *Intensity control*, *Frequency control*, *Masked activating sound*) on the P1/N1 amplitude, using R (55).

##### 5.2.2 Cerebral vestibular-evoked potentials

Peak amplitudes of evoked potentials were measured for each acoustic stimulus and body orientation separately. The peak-to-peak amplitude for biphasic components (e.g., Na/Pa or N*/P*) was calculated as the difference between the amplitudes of the two opposite peaks (e.g., N and P). When a component was not clearly identifiable, its peak value was set to 0.

Given the non-normal distribution of the data, an aligned rank transform (ART) ANOVA (56) was used to assess the main effects of *Sound* and *Orientation*, as well as their interaction, on the peak-to-peak amplitudes of each biphasic component. The ART ANOVA was conducted using the ARTool library in R (57), with *Sound* (*Activating sound*, *Intensity control*, *Frequency control*, *Masked activating sound*) and *Orientation* (*Upright sitting, Lying supine*) as fixed factors, and their interaction included in the model. Subjects were treated as a random factor. The model was specified as: *Amplitude ∼ Sound*Orientation + (1|Subject)*. Post-hoc multifactor contrasts tests were conducted following the aligned rank transform contrasts (ART-C) procedure (58), and *p*-values were adjusted using Holm’s correction.

#### 5.3 Assessing myogenic contributions to cervical vestibular-evoked myogenic potentials

To assess the contribution of muscle activity to vEPs, we computed cross-correlation coefficients between recordings from scalp, lIO, and rSCM electrodes. Cross-correlation was computed by temporally shifting one signal relative to the other and calculating their similarity at each time lag, allowing us to evaluate both the strength and latency of EEG–EMG coupling (59). For each EEG channel, the peak cross-correlation coefficient (*R*) and its corresponding time lag (in ms) were extracted to determine the temporal alignment between cerebral and myogenic signals. Cross-correlation analyses were performed on grand-average evoked responses elicited by the *Unmasked activating sound* in the upright sitting position, as this was the only posture in which muscle tone remained sufficient (60). In this context, residual VEMPs over the ipsilateral rSCM and contralateral lIO were plausible. EEG channels with *R* values exceeding the 95^th^ percentile were considered significantly associated to myogenic activity. A lag window of ±5 ms was defined as physiologically plausible considering conduction delay between myogenic signals propagating through muscle, skin, and other soft tissues to the scalp. This is consistent with typical surface EMG conduction velocities, which range from approximately 3−6 m/s and result in delays of a few milliseconds over short distances (61).

## RESULTS

### 1. Cervical vestibular-evoked myogenic potentials

As expected, both the *Unmasked* and *Masked activating sounds* (105 dB, 500 Hz) reliably evoked cVEMPs over the right SCM muscle, characterized by biphasic P1/N1 responses. In contrast, the *Frequency* and *Intensity control* stimuli did not elicit any consistent response (**Figure 2a**).

**Figure 2.**
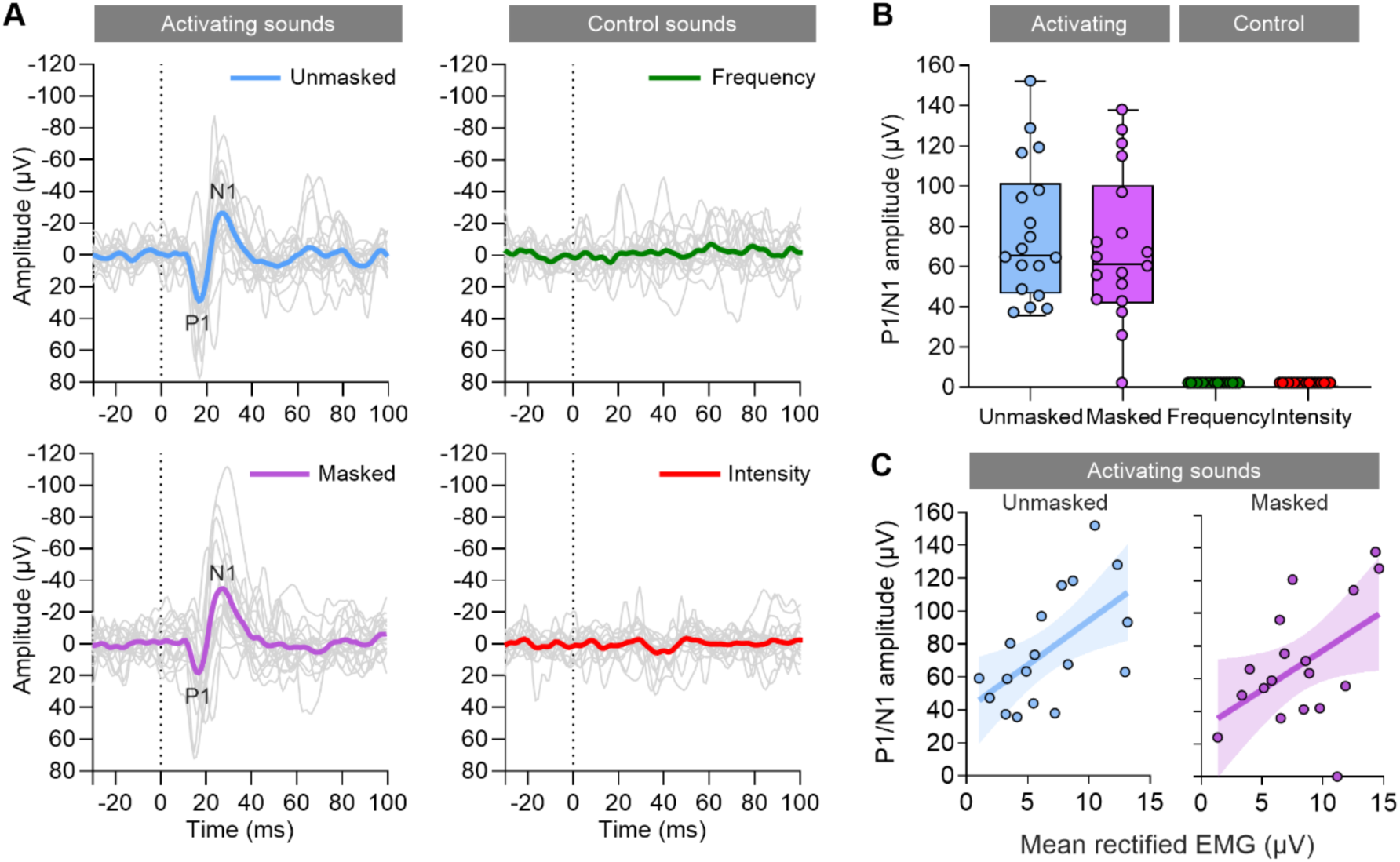
Cervical vestibular-evoked myogenic potentials recorded from the right sternocleidomastoid muscle. **(A)** Grand-average cervical vestibular-evoked myogenic potentials (thick colored lines; *n* = 18) and individual participant traces (thin grey lines) in response to two acoustic stimuli targeting the otolithic receptors (*Unmasked activating sound, Masked activating* sound) and two control stimuli matched for frequency (*Frequency control*) and intensity (*Intensity control*). Responses are time-locked to the onset of the tone pip (vertical dotted line). **(B)** Box-and-whisker plots showing P1/N1 peak-to-peak amplitudes across participants (*n* = 18) for each stimulus. Boxes indicate the interquartile range; horizontal lines within boxes represent the median; whiskers extend from the 5^th^ to the 95^th^ percentile. **(c)** Correlation between P1/N1 peak-to-peak amplitude and background EMG activity (mean rectified amplitude during the pre-stimulus baseline).

The P1/N1 peak-to-peak amplitude was significantly modulated by the type of acoustic stimulus (Friedman’s test: *χ*²(3) = 50, *p* < 0.001, Kendall’s *W* = 0.93). Post-hoc Dunn’s test revealed that both the *Unmasked activating sound* (mean ± SD: 73.3 ± 39.1 μV; *Mdn*: 64 μV) and the *Masked activating sound* (mean ± SD: 63.4 ± 33.9 μV; *Mdn*: 57 μV) elicited significantly larger P1/N1 amplitudes compared to the *Intensity* and *Frequency control* stimuli (*p* < 0.001). Importantly, masking the tone pip did not significantly change the P1/N1 amplitude, which did not differ between the *Unmasked* and *Masked activating sounds* (**Figure 2b**).

The cVEMPs amplitude exhibited substantial interindividual variability (range: 26−152 μV for the *Unmasked activating sound*; 24−136 μV for the *Masked activating sound*, with one participant showing no cVEMPs). This variability was in part related to differences in SCM contraction levels, as P1/N1 amplitudes correlated significantly with the mean rectified EMG recorded during the 30 ms preceding stimulus onset (*Activating sound*: *r*² = 0.34, *p* = 0.01; *Masked activating sound*: *r*² = 0.24, *p* = 0.04) (**Figure 2c**).

### 2. Cerebral vestibular-evoked potentials morphology

**Figure 3** illustrates the grand average waveforms elicited by the three unmasked tone pips, recorded in both the upright sitting and lying supine positions from several midline electrodes, as well as from the rSCM and lIO electrodes. Responses to the *Masked activating sound*, which present different morphologies, are illustrated separately (**Figure 5**).

**Figure 3.**
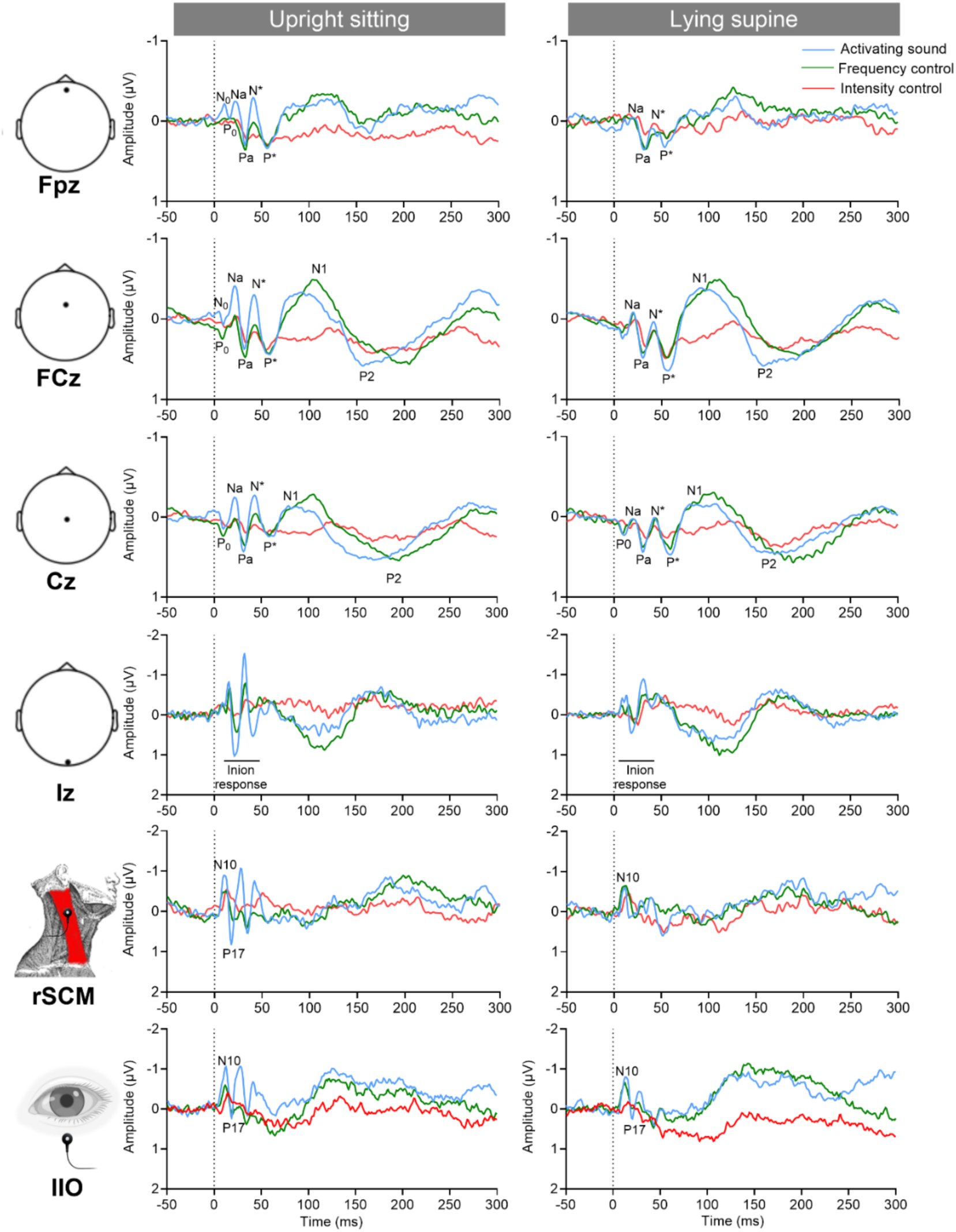
Vestibular-evoked potentials elicited by three unmasked acoustic stimuli. Grand-average waveforms (*n* = 18) of vestibular-evoked potentials in response to an unmasked acoustic stimulus targeting the otolithic receptors (*Activating sound*), and two control stimuli matched for frequency (*Frequency control*) and intensity (*Intensity control*). Recordings were obtained from four midline scalp electrodes (Fpz, FCz, Cz, Iz), the left infra-ocular electrode (lIO), and the right sternocleidomastoid muscle (rSCM), in both the upright sitting position and the lying supine position. Responses are time-locked to the onset of the tone pip (vertical dotted line).

Consistent with previous studies on auditory-evoked potentials and vEPs (62, 63), tone pips reliably elicited short- and middle-latency components at fronto-central electrodes (FCz, Cz). On electrode FCz, these include middle-latency components likely equivalent to N_0_ (e.g., peak latency for *Unmasked activating sound*, upright sitting [mean ± SD]: 7 ± 0.5 ms), P_0_ (peak latency: 10 ± 0.4 ms), Na (20 ± 0.4 ms), Pa (30 ± 0.4 ms), N* (41 ± 4.1 ms), and P* (54 ± 5.7 ms), as well as late components N1 (93 ± 22.8 ms) and P2 (160 ± 30.2 ms). A comparable middle-latency morphology was observed at frontopolar midline electrode Fpz, which also exhibits an early negative N_0_ component (8 ± 1.3 ms), selectively for the *Unmasked activating sound* (20, 64), especially in the upright sitting position.

Both *Unmasked activating sound* and *Frequency control* stimuli presented at 105 dB evoked an inion-type response at electrode Iz (60), which was more pronounced in the upright sitting position. Additionally, successive biphasic responses, including N10/P17 components, were recorded at electrodes rSCM and lIO in response to the *Unmasked activating sound* in both upright and sitting positions. Notably, the N10/P17 at electrode lIO was attenuated in the lying supine position, whereas only the N10 component persisted at rSCM in the lying supine position in response to both *Unmasked activating sound* and *Frequency control* stimuli. In contrast, electrodes Fpz, FCz, and Cz did not exhibit the early biphasic N10/P17 component. Instead, they displayed a different response morphology characterized by middle-latency components Na (e.g., peak latency for *Unmasked activating sound*, upright sitting: 21 ± 2.6 ms), Pa (30 ± 3 ms), N* (40 ± 3) and P* (52 ± 4.4 ms).

### 3. Influence of sound types and body orientation on cerebral vestibular-evoked potentials

The interaction between *Sound type* and *Body orientation* was assessed using a cluster-based permutation test within the 0–70 ms post-stimulus latency range, covering previously identified short- and middle-latency vEPs (20). This analysis did not include the *Masked activating sound* due to its altered vEPs morphology (see **Figure 5**), which made comparisons with other sound types impractical.

***Significant interactions between sound type and body orientation.*** The cluster-based permutation test revealed a significant interaction between *Sound type* and *Body orientation* factors within two distinct time windows associated with middle-latency components (**Figure 4a**). A first significant interaction (*p* < 0.05) was found in the 18–52 ms post-stimulus latency range, encompassing components Na, Pa, and N*. The interaction was pronounced over 32 electrodes, symmetrically distributed across the fronto-central scalp region, particularly around FCz electrode. Another significant interaction (*p* < 0.05) was found in the 50–70 ms post-stimulus latency range, primarily overlapping component P*. This interaction was pronounced over fronto-central electrodes with a left predominance, peaking at electrode FC5.

**Figure 4.**
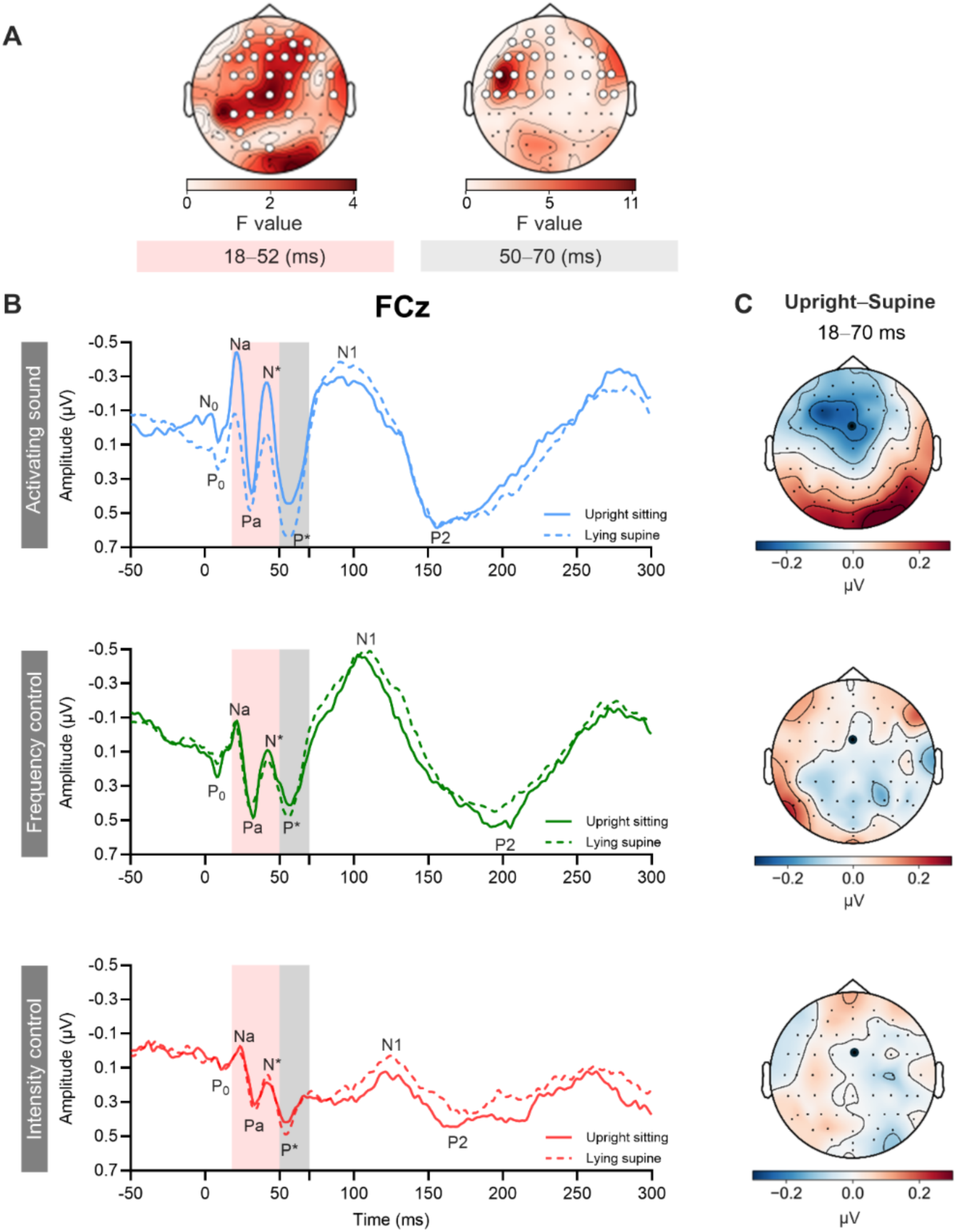
Results from the spatiotemporal cluster-based permutation analysis. (**A**) Averaged *F*-maps correspond to two post-stimulation latency windows: 18−52 ms and 50−70 ms after onset of the acoustic stimulation. Significant interactions between sound and body orientation for unmasked acoustic stimulations were identified using a cluster-based permutation test (32 electrodes for the 18−52 ms window; 27 electrodes for the 50-70 ms window; significant electrodes are highlighted in white). (**B**) Grand-average vestibular-evoked potentials (*n* = 18) recorded at FCz electrode in response to the acoustic stimulus targeting otolithic receptors (*Activating sound*) and two control stimuli (*Intensity control, Frequency control*), in both upright sitting and lying supine positions. Pink and grey shaded areas indicate time windows where a significant interaction between sound and body orientation was detected (18−52 ms and 50−70 ms, respectively). The 18−52 ms window includes middle latency components Na, Pa, and N*; the 50−70 ms window includes middle latency component P*. (**C**) Topographic maps showing the difference between upright sitting and supine positions for each acoustic stimulus, averaged over the 18–70 ms interval. Blue and red indicate negative and positive potential differences, respectively.

***Modulation of vEPs waveforms by body orientation.*** Grand-average vEPs waveforms recorded at electrode FCz for each acoustic stimulus and body orientation are shown in **Figure 4b**. In the 18–52 ms post-stimulus time window, two biphasic components were observed: Na/Pa (20−30 ms), and N*/P* (41−54 ms). In the 50–70 ms post-stimulus time window, a single monophasic component was identified, P* (54 ms).

The amplitude of Na/Pa and N*/P* was reduced in the lying supine position compared to the sitting upright position when the *Unmasked activating sound* was presented. In contrast, no modulation by body orientation was observed for both *Frequency control* and *Intensity control* stimuli.

***Topographic analysis of body orientation effects***. Averaged topographic maps illustrating the contrast between upright sitting and lying supine conditions in the 18−70 ms post-stimulus latency range were computed for each acoustic stimulus (**Figure 4c**). Consistent with the vEPs waveforms, an enhanced fronto-central negativity was observed for the *Unmasked activating sound*, whereas no clear topographic changes were evident for both control acoustic stimuli.

### 4. Effect of acoustic masking on cerebral vestibular-evoked potentials

To disentangle the components evoked by the masking noise from those evoked by the tone pip, we calculated the grand average response time-locked either to the tone pip onset or to the onset of the masking noise presented 50 ms earlier (**Figure 5**).

**Figure 5.**
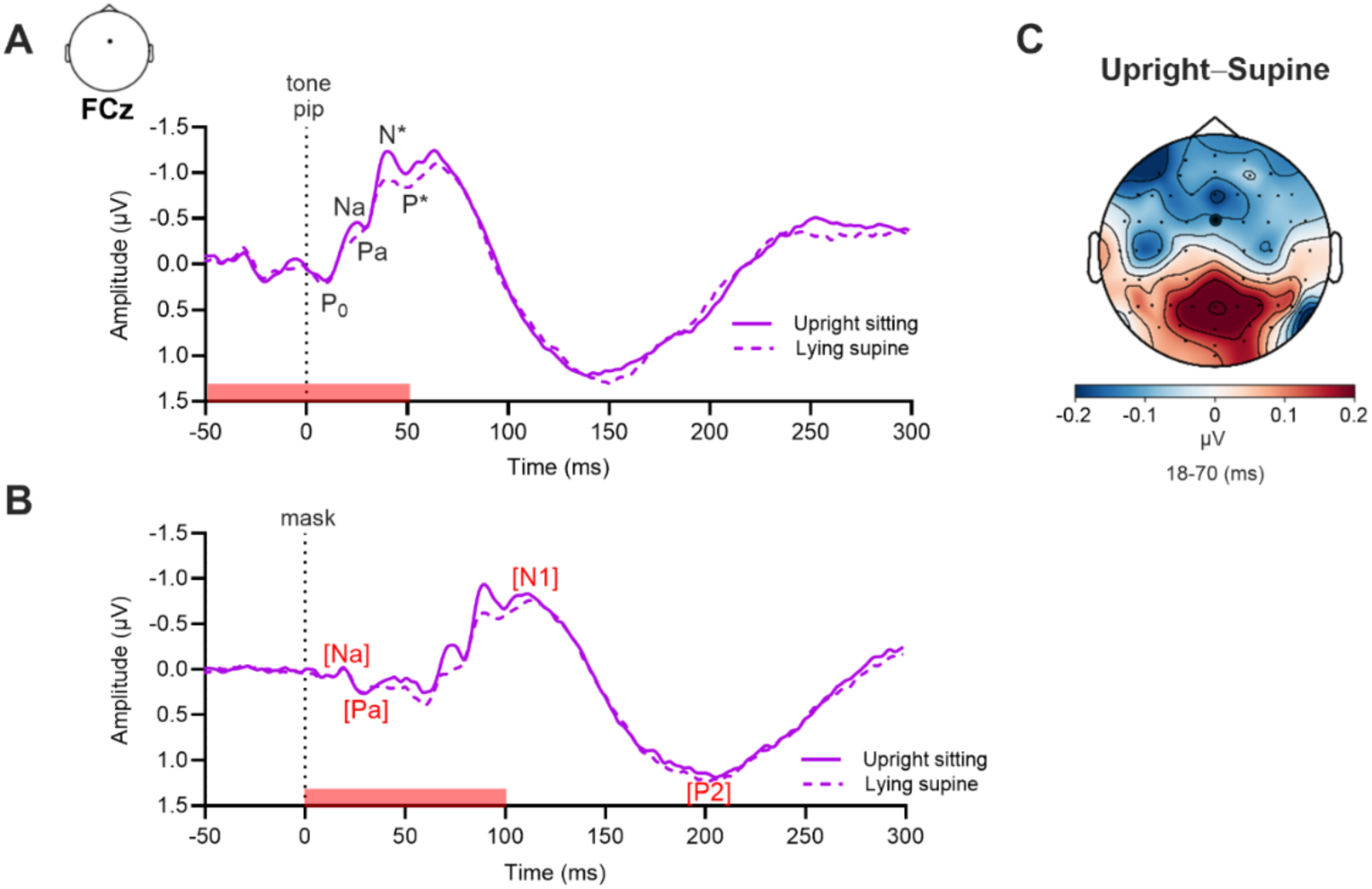
Vestibular-evoked potentials elicited by the masked activating sound. Grand-average vestibular-evoked potentials (*n* = 18) recorded at fronto-central electrode FCz in response to a masked acoustic stimulus targeting otolithic receptors (*Masked activating sound*), in both upright sitting (solid line) and lying supine (dashed line) positions. Responses are time-locked either to the onset of the tone pip (**A**) or to the onset of the masking noise (**B**). The duration of the masking noise (100 ms) is indicated by a red horizontal rectangle. **(C)** Topographic map of the difference between upright sitting and lying supine positions for the *Masked Activating sound*, time-locked to the onset of the tone pip and averaged over the 18−70 ms post-stimulation window. Blue and red indicate negative and positive potential differences, respectively.

Responses time-locked to the tone pip (**Figure 5a**) displayed markedly altered morphology compared to previous unmasked acoustic stimuli (compare **Figure 4**). The overall response amplitude was larger for the *Masked activating sound*, with short- and middle-latency components superimposed on a large biphasic deflection peaking at approximately 50 and 150 ms after tone pip onset. The following components could be identified: the short-latency P_0_ (peak latency: 10 ± 0.5 ms), and the middle-latency components Na (21 ± 2.8 ms), Pa (29 ± 3 ms), N* (40 ± 2.12 ms), and P* (51 ± 3.2 ms). Notably, only the Na/Pa and N*/P* components showed amplitude modulation by body orientation. In contrast to responses evoked by unmasked acoustic stimuli, no late auditory components (N1/P2) were observed.

Responses time-locked to the onset of the masking noise (**Figure 5b**) revealed that the large biphasic deflection described above corresponded to the late-latency auditory components N1 (peak latency: 92 ± 5.6 ms) and P2 (157 ± 7.2 ms), reflecting cortical processing of the masking noise itself. Early auditory components (Na/Pa), associated with initial auditory processing of the noise, were also observed. In contrast, there were no N*/P* components evoked by the masking noise, nor any indication that these components were modulated by body orientation, further supporting the interpretation that these middle-latency components reflect otolithic rather than auditory processing.

A topographic map of the contrast between upright sitting and lying supine was computed for the 18−70 ms post-stimulus time window, time-locked to tone pip onset (**Figure 5c**). Consistent with responses to the *Unmasked activating sound* (**Figure 4c**), an enhanced fronto-central negativity was observed in the upright position. This finding suggests that, despite the altered morphology induced by masking, the amplitude of middle-latency vEPs remained enhanced when participants were sitting upright.

### 5. Peak-to-peak amplitude analysis of middle-latency components

We conducted a peak-to-peak amplitude analysis at FCz electrode as (1) FCz middle-latency components have been identified as markers of otolithic information processing (20), and (2) our cluster-based permutation tests revealed a significant interaction between *Sound type* and *Body orientation* within time windows corresponding to these components over fronto-central scalp region. This approach allows for direct comparison across all acoustic stimuli, including the *Masked activating sound,* as peak-to-peak measurements remain valid regardless of waveform morphology. We analyzed the Na/Pa and N*/P* biphasic components for each sound type and body orientation (**Figure 6**).

**Figure 6.**
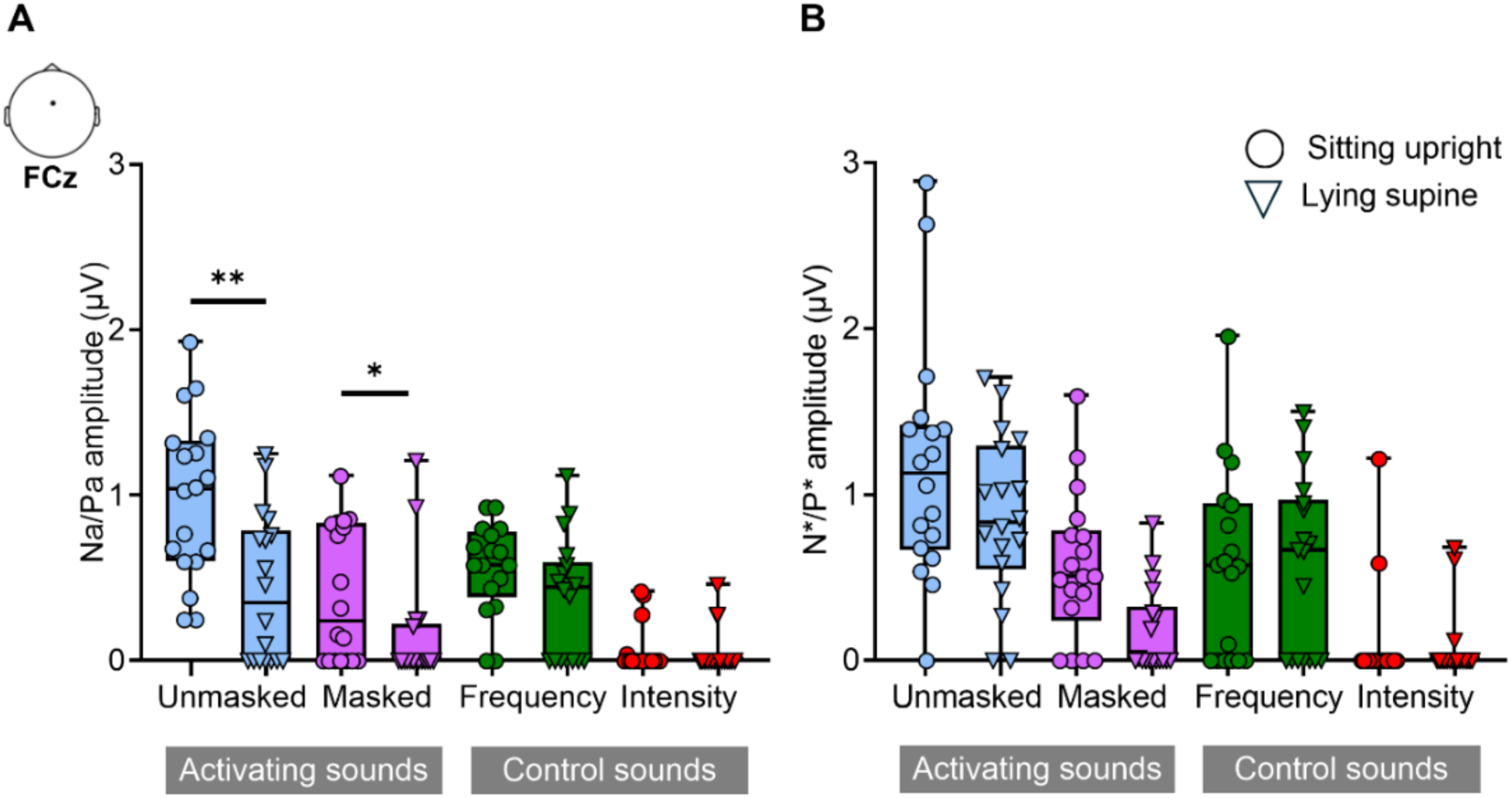
Peak-to-peak amplitudes of short- and middle-latency vestibular-evoked potentials at FCz electrode. Box-and-whisker plots showing the amplitudes of the biphasic responses Na/Pa (**A**) and N*/P* (**B**) elicited by each acoustic stimulus (*Unmasked and Masked activating sounds*, *Intensity and Frequency control sounds*), recorded at FCz electrode (*n* = 18). Boxes indicate the interquartile range; horizontal lines within boxes represent the median; whiskers extend from the 5^th^ to the 95^th^ percentile. Asterisks indicate significant differences between both body orientations (\**p* < 0.05, *\*\*p* < 0.01).

The ART ANOVA on Na/Pa peak-to-peak amplitudes revealed significant main effects of *Sound type* (*F*(*3, 119*) = 27.3, *p* < 0.001) and *Body orientation* (*F*(*1, 119*) = 22.1, *p* < 0.001), as well as a significant interaction between these factors (*F*(*3, 119*) = 5.5, *p* < 0.01). Post-hoc tests indicated that the Na/Pa amplitudes were significantly larger for the *Unmasked activating sound* (mean: 1.00 ± 0.6 μV; *Mdn*: 0.96 μV) compared to the *Masked activating sound* (mean: 0.28 ± 0.3 μV; *Mdn*: 0.10 μV; *p* < 0.001), *Frequency control* (mean: 0.55 ± 0.5 μV; *Mdn*: 0.59 μV; *p* < 0.001), and *Intensity control* (mean: 0.11 ± 0.3 μV; *Mdn*: 0.0 μV; *p* < 0.001). Post-hoc multifactor contrasts tests showed that Na/Pa amplitudes elicited by the *Unmasked activating sound* were significantly larger in the upright sitting position (mean: 0.98 ± 0.5 μV; *Mdn*: 1.04 μV) compared to the lying supine position (mean: 0.43 ± 0.4 μV; *Mdn*: 0.35 μV; *p* < 0.01). Similarly, Na/Pa amplitudes elicited by the *Masked activating sound* were significantly larger in the upright sitting position (mean: 0.39 ± 0.4 μV; *Mdn*: 0.23 μV) than in the lying supine position (mean: 0.10 ± 0.2 μV; *Mdn*: 0 μV; *p* < 0.05) (**Figure 6a**). By contrast, there was no significant difference in response amplitude between the upright sitting and lying supine positions for both control sounds (all *p*-values > 0.85).

The ART ANOVA on N*/P* peak-to-peak amplitudes showed a significant main effect of *Sound type* (*F*(*3, 119*) = 28.1, *p* < 0.001), but no significant main effect of *Body orientation* (*F*(*1, 119)* = 3.83, *p* = 0.052), nor interaction between factors (*F*(*3, 119*) = 1.9, *p* = 0.132) were found. Post-hoc tests indicated that the N*/P* amplitudes were significantly larger for the *Unmasked activating sound* (mean: 1.00 ± 0.6 μV; *Mdn*: 0.96 μV) compared to the *Masked activating sound* (mean: 0.28 ± 0.3 μV; *Mdn*: 0.10 μV; *p* < 0.001), *Frequency control* (mean: 0.55 ± 0.5 μV; *Mdn*: 0.59 μV; *p* < 0.001), and *Intensity control* (mean: 0.11 ± 0.3 μV; *Mdn*: 0.0 μV; *p* < 0.001). In addition, N*/P* amplitudes elicited by the *Frequency control* were significantly larger than those elicited by the *Masked activating sound* (*p* < 0.05) and *Intensity control* (*p* < 0.001). Finally, N*/P* amplitudes evoked by the *Masked activating sound* were significantly larger than those evoked by the *Intensity control* (*p* < 0.05) (**Figure 6b**).

### 6. Myogenic contributions to cerebral vestibular-evoked potentials

A clear N10/P17 biphasic component was observed at the inion (Iz electrode) in response to the *Unmasked activating sound*. Corresponding responses were also detected at rSCM and lIO EMG electrodes, with larger amplitudes in the upright position, and a marked attenuation in the supine position (**Figure 3**).

Cross-correlation analyses between EEG and EMG signals within the 0−70 ms post-stimulus time window revealed strong coupling, particularly between Iz and EMG channels (**Figure 7a−b**). Maximum cross-correlations were *R* = 0.84 at –1 ms delay for the Iz–rSCM pair, and *R* = 0.85 at –3 ms delay for Iz–lIO pair, indicating that EMG signals from rSCM and lIO preceded the EEG signal at Iz by approximately 1 ms and 3 ms, respectively. Only five scalp electrodes (Fp1, FC6, PO7, O1, and Iz) exceeded the 95^th^ percentile threshold (**Figure 7c**), indicating significant EEG–EMG coupling, thus reflecting a contribution of myogenic activity to the recorded signals at these sites.

**Figure 7.**
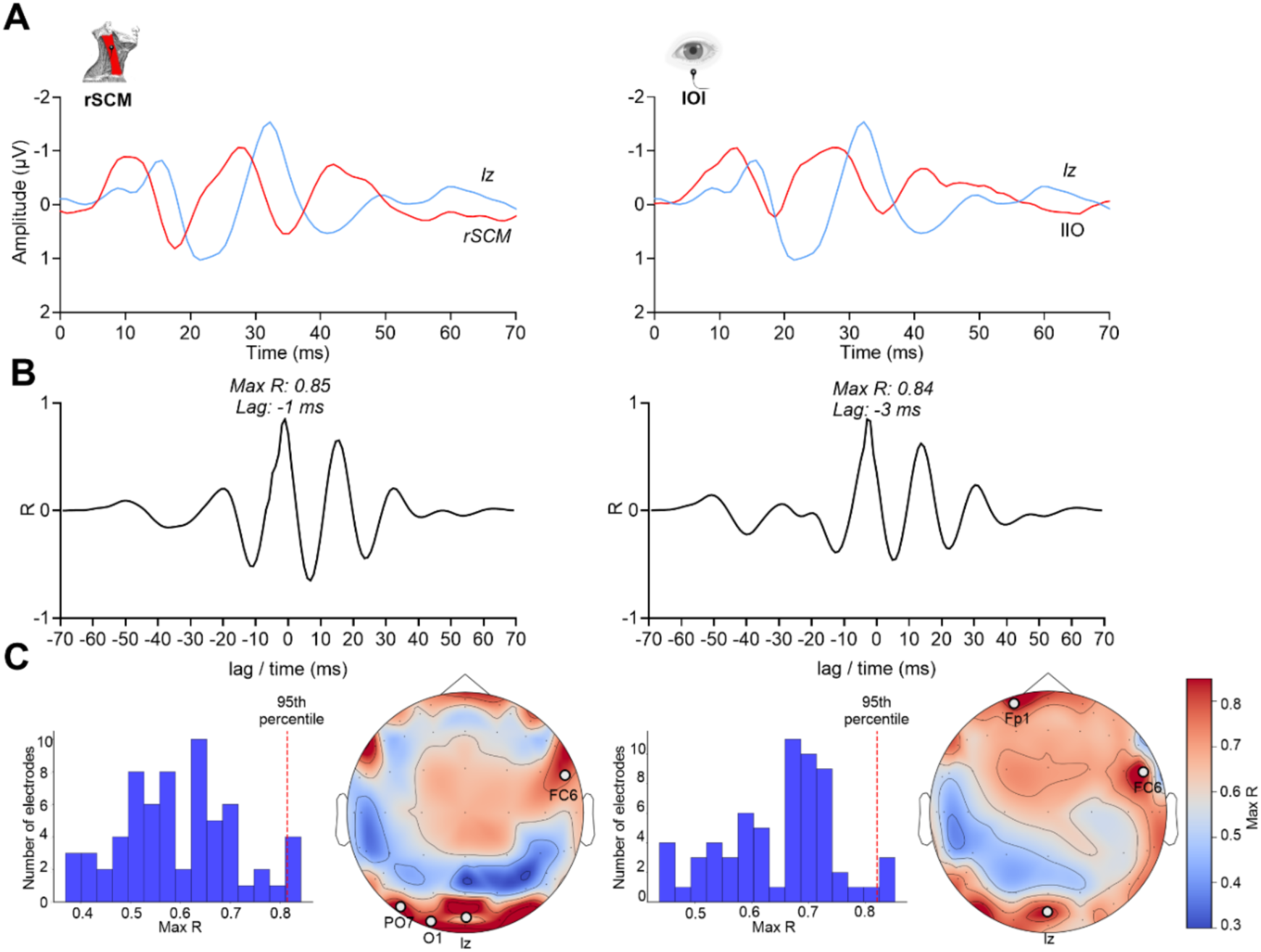
Cross-correlation between vestibular-evoked signals recorded from the right sternocleidomastoid (rSCM) and left inferior oblique (lIO) muscles in the upright sitting position. (**A**) Grand-average evoked responses from Iz (blue), rSCM (red on left panel) and lIO (red on right panel) electrodes (*n* = 18). (**B**) Cross-correlation functions between EEG and rSCM (left) and between EEG and lIO (right) calculated over the 0–70 ms post-stimulus time window. Peak cross-correlation values and their associated delays are reported; negative lags indicate that the EMG signal precedes the EEG signal. (**C**) Distributions of maximum cross-correlation (max *R*) values across 64 EEG electrodes. Red dashed lines indicate the 95^th^ percentile thresholds used to identify electrodes with significantly high correlation values. Scalp topographies showing the spatial distribution of maximum cross-correlation (max *R*). Warm colors indicate stronger correlations; cooler colors indicate weaker correlations. Fp1, FC6, PO7, O1, and Iz electrodes exhibit maximum *R* values above the 95^th^ percentile (M*ax R* > 0.80). Electrodes labeled on the maps showed peak *R* values with lag times between –3 ms and +3 ms.

Scalp topographies (**Figure 7c**) showed that these effects were most pronounced over posterior and fronto-lateral regions. Importantly, fronto-central EEG electrodes where significant modulations of vEPs amplitude by body orientation were elicited did not exhibit significant cross-correlation with EMG signals.

Finally, we assessed whether peak-to-peak amplitudes of Na/Pa and N*/P* at FCz electrode correlated with background EMG activity (mean rectified EMG over the 30 ms preceding stimulus onset) in the upright sitting position. No significant correlation was found between middle-latency responses and background EMG from rSCM (Na/Pa: *R* = 0.06, *p* = 0.35; N*/P*: *R* = 0.03, *p* = 0.51) or lIO (Na/Pa: *R* = 0.02, *p* = 0.54; N*/P*: *R* = 0.01, *p* = 0.61). Likewise, the inion response amplitude at Iz did not correlate with EMG activity from either muscle (rSCM: *R* = 0.10, *p* = 0.21; lIO: *R* = 0.06, *p* = 0.35).

## DISCUSSION

Our results indicate that acoustic vEPs provide a reliable physiological marker of otolithic processing. A key novel finding of this study is the identification of middle-latency components (Na/Pa and N*/P*), whose amplitudes are selectively modulated by body orientation relative to gravity, but only for acoustic stimuli activating otolithic receptors, and not for control sounds matched in frequency and intensity. This demonstrates that the dynamics of otolithic processing is influenced by factors such as head orientation and postural constraints within the gravitational field.

### 1. Validation of otolithic activation via cervical vestibular-evoked myogenic potentials

We first electrophysiologically validated the distinction between vestibular-activating sounds and control sounds. Activating sounds (105 dB, 500 Hz) reliably elicited classical biphasic P1/N1 cVEMPs responses over the SCM ipsilateral to the stimulus, consistent with prior studies (1, 23, 48). As expected, cVEMPs amplitude correlated with SCM background EMG level (23, 65).

Importantly, frequency- and intensity-matched control sounds failed to elicit cVEMPs, demonstrating their suitability as auditory controls in our EEG paradigm. We also developed a novel masked activating sound that reduced auditory perception and processing while preserving otolithic activation. The P1/N1 response to this masked sound was indistinguishable in amplitude from that evoked by the unmasked sound of the same intensity, confirming robust otolithic pathway activation.

Animal studies have shown that SVS predominantly activates saccular and utricular maculae (18) with negligeable semicircular canals contributions, except at higher sound pressure levels (reviewed in (66)). In conclusion, our stimuli reliably activate otolithic pathways, which project to vestibulo-collic, vestibulo-ocular, vestibulo-cerebellar, and vestibulo-thalamo-cortical targets (1, 31, 32).

### 2. Spatiotemporal dynamics of vestibular-evoked cortical potentials

Our study advances the understanding of vestibular information processing by examining the combined effects of acoustic stimulation and body orientation on cortical vEPs. The scalp EEG components we recorded replicated key features of cortical vEPs reported in prior non-invasive (20, 22, 60) and recent intracranial (21, 22) EEG studies. These included an early P10/N17 complex at posterior electrodes, followed by prominent Na/Pa (20–30 ms) and N*/P* (41–54 ms) components over fronto-central regions.

#### 2.1. Short-latency components (P10/N17)

The earliest responses likely reflect a mix of myogenic and neurogenic contributions. Loud sounds can evoke inion responses compatible with muscular origins (67, 68). Such myogenic activity takes the form of a VEMP with a reversed waveform polarity, as it probably originates from the SCM and trapezius muscles insertions (reviewed in (63)). Extraocular muscle activation may also contribute to early responses near Fpz or Pz electrodes (64). However, neurogenic contributions to these short-latency components are also supported by evidence of responses from the brainstem, cerebellum (64, 36, 60), and multisensory vestibular cortex (36, 69). Intraoperative vestibular nerve stimulation elicits cortical responses as early as 6 ms (69), and intracranial EEG recordings show short-latency activation in the Heschl’s gyrus and posterior insula around 10–20 ms (21). Similar short-latency vEPs have been recorded in non-human animals cerebral cortex following direct vestibular nerve stimulation (70–72).

#### 2.2. Middle-latency components (Na/Pa and N*/P*)

The most robust vestibular-evoked responses in our dataset were Na/Pa (20–30 ms) and N*/P* (41–54 ms) complexes at FCz electrode. These responses align with prior reports linking them to sources in the superior temporal gyrus (63, 73, 74). These middle-latency responses could reflect mixed auditory/vestibular or neck/scalp muscles activity (reviewed in (63)). Yet, a neurogenic vestibular origin is strongly supported by the fact that the N*/P* component was absent in a patient with a vestibular loss (20), and our cluster-based permutation analysis showed amplitude modulation by body orientation, specific to otolith-activating sounds.

Notably, our results further suggest that Na/Pa is also sensitive to otolithic processing, an effect not previously reported (20). This aligns with intracranial EEG studies showing SVS-evoked responses in the posterior insula and parietal operculum at 20 ms and 30 ms (22), compatible with our Na/Pa latency. Together, these observations indicate that middle-latency vEP components index early cortical processing of vestibular signals, with minimal muscular contamination.

### 3. Influence of body orientation on middle-latency vestibular-evoked potentials

Our results revealed a consistent reduction of middle-latency vEPs as a function of body orientation. Cluster-based permutation analysis showed a significant interaction between sound type and posture during the Na/Pa and N*/P* time window over fronto-central regions. Peak-to-peak analyses at FCz electrode confirmed that this interaction was significant for Na/Pa only, not for N*/P*. Importantly, Na/Pa amplitude was significantly reduced in the supine vs. upright position for both masked and unmasked activating sounds, with no modulation for control sounds. This pattern strongly suggests that the observed reduction reflects changes in otolithic, not auditory, processing. Several non-exclusive mechanisms may account for the attenuated middle-latency vEPs in the supine condition.

#### 3.1. Peripheral mechanisms

One possible explanation is altered otolithic receptor sensitivity when changing orientation in the pitch plane. One might argue that in the upright position, utricular and saccular hair cells are optimally oriented for detecting linear accelerations (75, 76), enhancing afferent discharge. However, while electrophysiological recordings in non-human primates indicate that the firing discharge of primary afferents strongly relates to the angle of static tilt in the pitch plane (38), some afferents show drastic decrease in firing rate in the supine position, while others show the opposite pattern (39, 40). In addition, SVS activate otolithic receptors in a non-physiological way which cannot be compared to classical mechanoelectrical transduction of gravito-inertial forces by the otolithic maculae. It has been proposed that SVS creates fluid pressure waves that displace directly hair bundles from otolithic receptors, “bypassing the mechanics of the otolithic maculae and even the otoconial layer” (17). In the supine position, the sustained tilt of hair bundles in otoconial membrane may reduce responsiveness to SVS-induced pressure waves, or limit their range of motion due to mechanical constraints.

Another potential mechanism involves changes in intralabyrinthine and intracranial fluid pressure in the supine position (77). There is mixed evidence regarding the influence of intracranial pressure: some studies report reduced oVEMPs in tilted positions (77), while others find no effect (78–80). Critically, changes in intracranial pressure do not affect typical auditory responses, such as transient evoked otoacoustic emissions (81, 82), high-frequency tympanometry (83), and modestly influence distortion-product otoacoustic emissions (84). This aligns with our observation of unchanged EPs amplitude for control acoustic stimuli in the supine position.

#### 3.2. Central mechanisms

The reduction in vEPs in the supine position more likely reflects central modulation of vestibular processing. Multisensory perception is profoundly influenced by orientation relative to gravity (85), and vestibular processing is further modulated by postural constraints, cognitive factors, and predictive coding (86).

In our study, the reduction in vEPs suggests a context-dependent down-weighting of otolithic signals when their relevance for postural control is reduced. While vestibular signals are critical for balance in the upright position (87), the minimal postural demand in the supine position may lead to central down-weighting of otolithic signals, while auditory processing remains unaffected. This relates to the selective reduction in middle-latency vEPs (Na/Pa and N*/P*), with preserved late auditory components (N1/P2) in our study. Supporting this hypothesis, cVEMPs and oVEMPs amplitudes increase under elevated postural threat, such as standing on elevated platforms, compared to recordings at ground level (88, 89). Additionally, cognitive load modulates cVEMPs amplitude (90), suggesting a complex interplay between attention, posture, and vestibular processing.

While the exact pathways through which postural constraints and predictive coding mechanisms operate remain unclear, several lines of evidence suggest plausible substrates. Recent studies have revealed decreased release of excitatory neurotransmitter in the parieto-insular vestibular cortex during attentional tasks (91). A similar mechanism could contribute to attenuate vEPs in the supine position, where postural demand is minimal. The cerebellum, which is richly interconnected with the vestibular nuclei, plays a pivotal role in the “dynamic prediction of the sensory consequences of gravity to ensure postural and perceptual stability” (92, 45). It may modulate multisensory integration within vestibular nuclei, thereby reducing ascending input to the multisensory vestibular cortex and contributing to the observed reduction in vEPs amplitude. While the efferent vestibular system could theoretically modulate synaptic transmission between otolithic receptors and afferent fibers, its capacity for short-term modulation of afferent discharge appears limited in mammals (93). Furthermore, neither predictive coding mechanisms related to active self-motion nor neck proprioceptive signals modulate the firing rate of vestibular afferents (94), making it unlikely that this system accounts for the rapid, posture-dependent changes in vEPs observed in our study.

### 4. Influence of auditory masking on vestibular-evoked potentials

The masking procedure, classically used to suppress auditory perception (49, 50), altered the EEG response morphology. When time-locked to the masking noise, a large biphasic response N1/P2 (∼105–202 ms) reflected auditory processing, while time-locking to the tone pip preserved short- and middle-latency components (P_0_, Na/Pa, N*/P*), confirming vestibular activation despite auditory masking.

A critical dissociation emerged between auditory and vestibular responses: Late N1/P2 components typically associated with cortical auditory processing (95) were absent when time-locked to the tone pip but present when time-locked to the masking noise. In contrast, middle-latency components Na/Pa and N*/P* were not evoked by the masking noise alone and exhibited modulation by body orientation only when time-locked to the tone pip. This dissociation supports the otolithic contribution to middle-latency vEPs.

Furthermore, the topographic distribution of the masking condition revealed a fronto-central negativity enhanced in the upright posture, mirroring the *Unmasked activating sound*. This orientation-dependent modulation, preserved even with auditory masking, highlights the specificity of middle-latency vEPs for otolithic signals and their sensitivity to body orientation.

### 5. Neurogenic and myogenic contributions to vestibular-evoked potentials

Cross-correlation analyses revealed that EMG activity from rSCM and lIO muscles preceded EEG responses at posterior channels (Iz) by 1 to 3 ms, indicating a myogenic contribution for early components such as P10/N17. This aligns with previous studies identifying these responses as part of the inion response (60). In contrast, no EMG–EEG coupling was observed at fronto-central electrodes, reinforcing the interpretation that Na/Pa and N*/P* reflect cortical vestibular processing rather than myogenic contamination. Their distinct topography and timing further support a neurogenic origin for these middle-latency responses, confirmed by recent intracranial EEG recordings (21, 22).

Overall, our results highlight regional specificity: Posterior and frontal electrodes (e.g., Iz, Fp1) are prone to myogenic contamination, whereas midline frontocentral components (Na/Pa, N*/P*) likely originate from cerebral vestibular networks. This functional dissociation is critical for interpreting vEPs origins and identifying reliable cerebral markers of otolithic processing.

### 6. Conclusions

Our findings establish middle-latency vEPs components Na/Pa and N*/P* as robust markers of otolithic information processing. Their sensitivity to body orientation and acoustic specificity suggests they reflect multisensory integration within the vestibulo-thalamo-cortical and cerebellar networks.

This work advances our understanding of how the brain encodes gravity-sensitive signals, with implications for higher-order vestibular functions. The orientation-dependent reduction in vEPs aligns with extensive evidence that tilted body orientation alters multisensory perception, affecting verticality perception (85), body representation (96), tactile processing (97), and spatial navigation (98).

Finally, our results have direct implications for neuroimaging studies: functional MRI studies investigating SVS or galvanic vestibular stimulation all have been conducted with participants in a supine position. Given our demonstration of reduced vestibular responses in the supine position, functional MRI investigations of vestibular brain networks and spatial navigation systems must account for posture-dependent modulation of brain responses more systematically.

## ACKNOWLEDGMENTS

We thank Dr. Olivier Macherey for the calibration of the acoustic stimuli, and Dr. Laure Spieser and Dr Anne-Sophie Dubarry for guidance in EEG data analysis. We are grateful to Dany Paleresompoulle and Pierre Cauvin for their technical support.

## GRANTS

This study was supported by the ANR VESTISELF project, grant ANR-19-CE37-0027 of the French Agence Nationale de la Recherche (to CL).

## DISCLOSURES

Authors report no conflict of interest.

## AUTHOR CONTRIBUTIONS

Y.B., Y.C., and C.L. conceived and designed research;

P.K., L.S., and Y.B. performed experiments;

P.K., L.S., Y.B., and C.L. analyzed data;

P.K., L.S., Y.B., Y.C., and C.L. interpreted results of experiments;

P.K., L.S., Y.B., and C.L. prepared figures;

P.K., L.S., and C.L. drafted manuscript;

P.K., L.S., Y.B., Y.C., and C.L. edited and revised manuscript;

P.K., L.S., Y.B., Y.C., and C.L. approved final version of manuscript.

## Notes

### Competing Interest Statement

The authors have declared no competing interest.

